# Dysregulated RASGRP1 expression through RUNX1 mediated transcription promotes autoimmunity

**DOI:** 10.1101/656454

**Authors:** Matthijs Baars, Thera Douma, Dimitre R. Simeonov, Darienne R. Myers, Saikat Banerjee, Kayla Kulhanek, Susan Zwakenberg, Marijke P. Baltissen, Sytze de Roock, Femke van Wijk, Michiel Vermeulen, Alexander Marson, Jeroen P. Roose, Yvonne Vercoulen

## Abstract

RasGRP1 is a Ras guanine nucleotide exchange factor, and an essential regulator of lymphocyte receptor signaling. In mice, *Rasgrp1* deletion results in defective T lymphocyte development. *RASGRP1*-deficient patients suffer from immune deficiency, and the *RASGRP1* gene has been linked to autoimmunity. However, how RasGRP1 levels are regulated, and if RasGRP1 dosage alterations contribute to autoimmunity remains unknown. We demonstrate that diminished Rasgrp1 expression caused defective T lymphocyte selection in C57BL/6 mice, and that the severity of inflammatory disease inversely correlates with Rasgrp1 expression levels. In patients with autoimmunity, active inflammation correlated with decreased *RASGRP1* levels in CD4^+^ T cells. By analyzing H3K27 acetylation profiles in human T cells, we identified a *RASGRP1* enhancer that harbors autoimmunity-associated SNPs. CRISPR-Cas9 disruption of this enhancer caused lower RasGRP1 expression, and decreased binding of RUNX1 and CBFB transcription factors. Analyzing patients with autoimmunity, we detected reduced RUNX1 expression in CD4^+^ T cells. Lastly, we mechanistically link RUNX1 to transcriptional regulation of *RASGRP1* to reveal a key circuit regulating RasGRP1 expression, which is vital to prevent inflammatory disease.

## Introduction

RasGRP1, a Ras guanine nucleotide exchange factor (RasGEF), is a key protein regulating effector kinases upon T cell receptor (TCR) signaling[1]. RasGRP1 expression levels regulate the output of lymphocyte receptor signaling in a dose-dependent manner [2, 3]. Complete lack of *Rasgrp1* in mice results in a defect in positive selection of thymocytes (developing T cells), and immune deficiency, both in wildtype mice and mice expressing a transgenic TCR [4, 5]. Recent studies described *RASGRP1* loss in patients with immune deficiencies [6, 7], and RasGRP1 loss-of-function in autoimmune lymphoproliferative syndrome (ALPS) [8].

Dysregulation of RasGRP1 expression levels has been suggested to play a role as well in leukemia and autoimmunity. Increased expression of Rasgrp1 has been detected in murine models and patients with T-ALL [9, 10], and some studies demonstrated aberrant expression levels of RasGRP1 in patients with autoimmunity. For example, patients with systemic lupus erythematosus with splice variants of RasGRP1 expressed decreased RasGRP1 protein levels [11], while in rheumatoid arthritis increased mRNA and a contrasting decrease in RasGRP1 protein was shown in total T cells [12]. Furthermore, single nucleotide polymorphisms (SNPs) in the *RASGRP1* locus have been linked to autoimmunity [13–15]. For instance, SNPs in coding regions have been shown to affect lymphocyte receptor signaling in cell lines [2], and in mice [16]. It has not been established if non-coding SNPs in *RasGRP1* affect its expression levels. In sum, it is unknown how RasGRP1 expression is regulated in T cells and if RasGRP1 dosage alterations may impact T cell function and immunological health. We set out to mechanistically understand how RasGRP1 expression levels are regulated and investigate if aberrant expression of RasGRP1 may contribute to inflammatory disease.

## Results

To test whether decreased Rasgrp1 expression levels can in principal cause autoimmunity, we assessed immune-phenotypes of WT (+/+) to *Rasgrp1*-heterozygous (+/−) and *Rasgrp1*-KO (−/−) mice. All mice were on a C57BL/6 background, which is not prone to develop autoimmunity [17]. Thymocytes of *Rasgrp1* heterozygous mice express approximately half of normal Rasgrp1 protein levels (Fig 1A). Analysis of stages of thymocyte development (Fig 1B) by flow cytometry showed that aberrant Rasgrp1 expression impaired positive thymocyte selection. As reported previously [18], we observed that *Rasgrp1*^−/−^ mice displayed an accumulation of CD44^−^CD25^+^ double negative (DN3) thymocytes, indicating defective β-selection. By contrast, *Rasgrp1*^+/−^ mice did not show accumulation of this early T cell progenitor population directly prior to β-selection (Fig 1C, Fig 1-Suppl A). The positive selection process shapes the T cell receptor (TCR) repertoire as assembled, mature TCRs are functionally tested in this selection process [19] and alterations in the repertoire can lead to self-recognition in the periphery and autoimmune diseases. We observed that intact Rasgrp1 expression is critical for positive selection; the numbers of TCRβ^+^CD69^+^ double positive (DP) thymocytes and CD4^+^ SP cells were decreased in *Rasgrp1*^+/−^ mice (Fig 1D,E, Fig 1-Suppl B-C). As expected and reported [18], without any Rasgrp1 expression (in *Rasgrp1*^−/−^ mice) these two populations were nearly absent (Fig 1D,E, Fig 1-Suppl B-C). These results showed an inverse effect of RasGRP1 expression on thymic positive selection; half the RasGRP1 protein dosage already resulted in less efficient positive selection.

**Figure 1.**
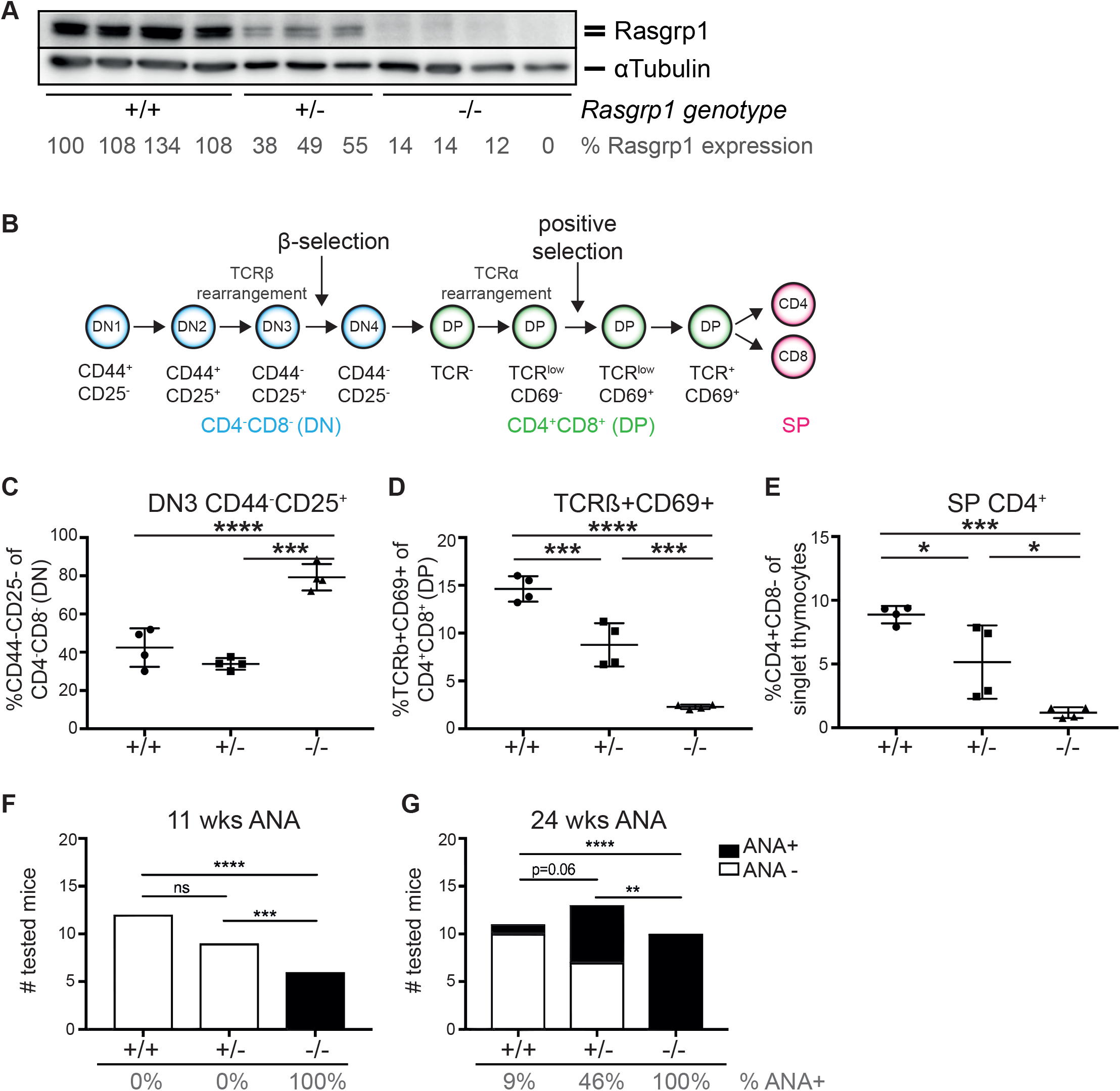
Limited RasGRP1 expression results in dose-dependent autoimmunity in mice. (A) Western blot of RasGRP1 protein levels expressed by thymocytes of 11 weeks old mice with different genotypes: +/+ (WT), +/− (*Rasgrp1* heterozygote), −/− (*Rasgrp1* KO). (B) Schematic representation of thymocyte positive selection. (C-E)Thymocytes were isolated from 5 weeks old mice and different stages of thymocyte selection were analyzed by flow cytometry. Shown are average percentages ± S.D. of (C) CD44-CD25^+^ (DN3) of gated CD4^−^CD8^−^ cells (double negative, DN), (D) TCRß^+^CD69^+^ of CD4^+^CD8^+^ (double positive, DP) cells, and (E) CD4^+^CD8^−^ (CD4 single positive, SP) of total thymocytes. One-way ANOVA was used. (F,G) Anti-nuclear antibody presence (ANA) was determined in sera isolated from mice at 11 weeks old (E, N=6, N=9, N=6), and 24 weeks old (F, N=7, N=13, N=10). Bars depict numbers of mice tested positive (black), and negative (white), below is the percentage of ANA positive samples. Fisher’s exact test was used. *p<0.05, ** p<0.01, ***p<0.001, ****p<0.0001.

Next, we analyzed serum for the presence of anti-nuclear antibodies, a commonly used indicator for autoimmune features (ANA, Fig 1-suppl D). In 11 weeks old mice, we observed that absence of Rasgrp1 led to ANA production, while *Rasgrp1*^+/−^ and WT mice produced no ANA at all (Fig 1F). These results agreed with earlier studies performed in *Rasgrp1*^−/−^ mice [20, 21]. However, by age 24 weeks we also detected serum ANA in approximately half of the heterozygous *Rasgrp1*^+/−^ mice, while only 1 out of 11 wildtype mice were ANA positive (Fig 1G). Thus, decreased expression of Rasgrp1 led to an increased risk of development of autoimmunity in C57BL/6 mice over time.

Based on these *in vivo* findings in C57BL/6 mice, we hypothesized that RasGRP1 expression could be dysregulated in human autoimmunity. Previous reports showed that peripheral blood total T cells (Both CD4^+^ and CD8^+^) from patients with Rheumatoid Arthritis (RA) display higher *RASGRP1* mRNA levels than healthy donors, and the expression levels of *RASGRP1* inversely correlate with disease activity scores [12]. We set out to explore *RASGRP1* expression in patients with oligoarticular Juvenile Idiopathic Arthritis (JIA), a relapsing/remitting form of autoimmunity in children. We analyzed *RASGRP1* expression in peripheral blood CD4^+^ T cells from patients who were either in remission or showed active inflammation, and compared this to healthy controls. Furthermore, we analyzed CD4^+^ T cells from the site of active inflammation; synovial fluid (SF). We observed that *RASGRP1* mRNA expression was significantly decreased in CD4^+^ T cells from patients with active autoimmune disease compared to healthy controls, and this was most pronounced in synovial fluid at the site of active inflammation (Figs 2A, 2-SupplA,B). These results imply that there is active maintenance of *RASGRP1* expression levels in CD4^+^ T cells with instances of decreased expression of *RASGRP1* mRNA levels specifically under autoimmune inflammatory conditions and motivated us to next investigate regulation of *RasGRP1* mRNA expression.

**Figure 2.**
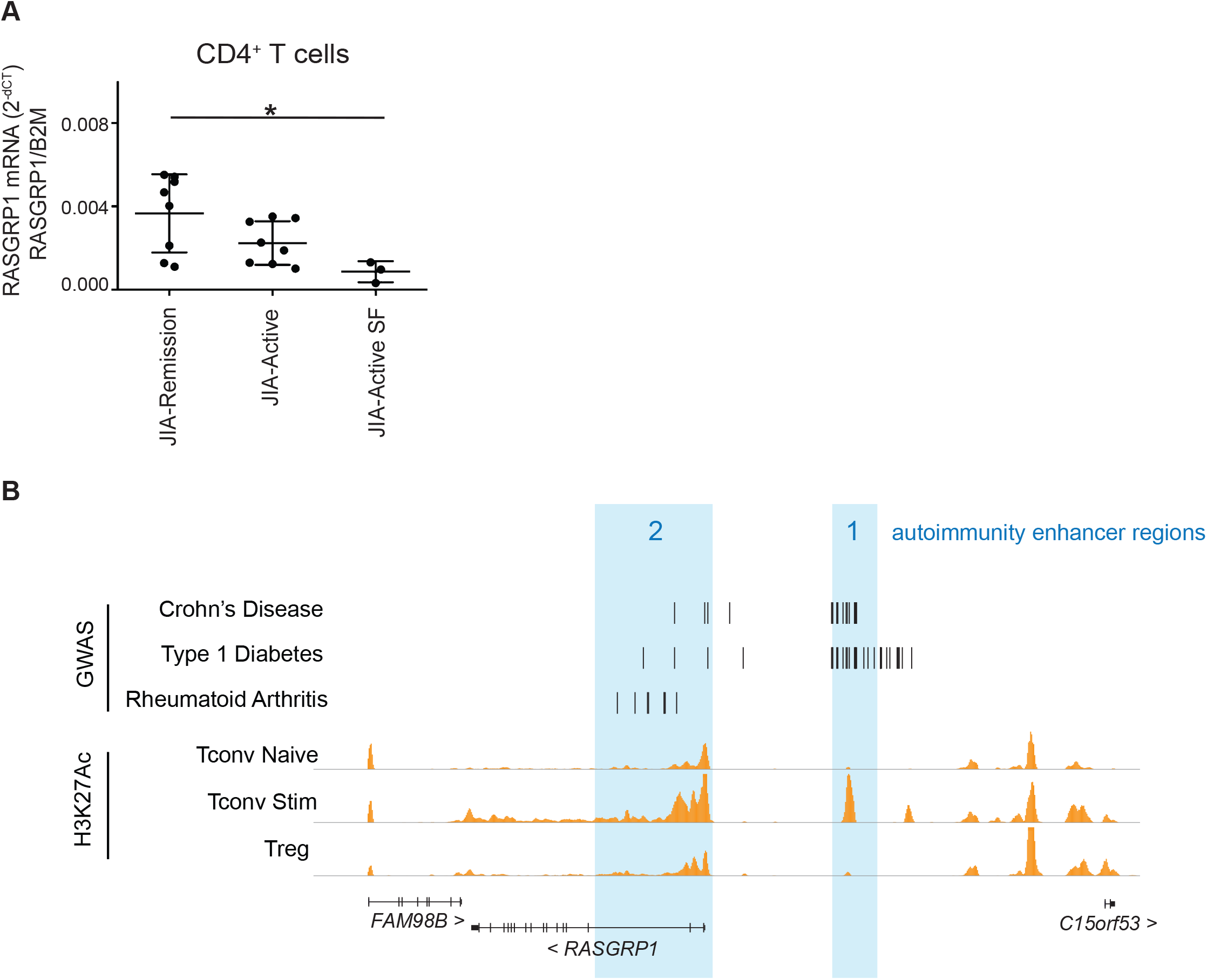
RASGRP1 levels and expression regulation are associated with human autoimmunity. (A) RASGRP1 mRNA levels in CD4+ T cells were isolated from juvenile idiopathic arthritis patients during disease remission blood (JIA-Remission, N=8), active inflammatory disease blood (JIA-Active, N=8), and synovial fluid (JIA-Active SF, N=3). Shown are averages ± S.D. 2^−dCT^ values of RasGRP1 corrected for B2M. One-way Anova, with Holm-Sidak’s multiple comparisons test was used for statistical analysis. (B) RasGRP1 locus showing fine mapped disease-associated SNPs for Crohn’s Disease (CD)[14], Type 1 Diabetes (T1D)[15], and Rheumatoid Arthritis (RA)[13]. The GWAS data was overlapped with histone 3 lysine 27 acetylation (H3K27Ac) tracks for human primary T cells (Roadmap Epigenomics Project [40]): Naïve conventional T cells (Tconv Naïve), anti-CD3/anti-CD28 stimulated conventional T cells (Tconv Stim), and CD4^+^FOXP3^+^ regulatory T cells (Treg). 2 regions with H3K27 signal and colocalizing autoimmune SNPs are highlighted as autoimmunity-associated enhancer regions.

To identify non-coding elements that could affect *RASGRP1* gene expression in the context of autoimmunity, we analyzed the locations of candidate causal non-coding SNPs associated with autoimmunity. We found two distinct regions with clusters of candidate causal SNPs associated with Crohn’s disease, Type 1 Diabetes, and RA. One region was positioned upstream of the *RASGRP1* promoter, suggesting the presence of regulatory elements relevant to autoimmunity (Figs 2B, 2-Suppl B). Next, we analyzed Histone 3 lysine 27 acetylation (H3K27ac), indicating transcriptional enhancers. Enhancers are essential to tune gene regulation, and contribute to cell identity and function in health and disease [22, 23]. In human CD4^+^ T cells, the H3K27ac signal on *RasGRP1* elements was more pronounced in activated/stimulated T cells, and co-localized with the SNP clusters (Figs 2B). Notably, we identified a prominent, single H3K27ac peak marking a putative enhancer (autoimmunity-associated enhancer 1). These data suggest that *RASGRP1* expression in CD4^+^ T cells is regulated by enhancers and that SNPs in these elements could contribute to RASGRP1 dysregulation and autoimmunity.

To test whether the putative autoimmunity-associated enhancer 1 indeed regulates RASGRP1 expression, we perturbed this non-coding element with CRISPR-Cas9 editing in the Jurkat T cell lymphoma line [24]. First, we transfected cells with Cas9-Ribonucleoprotein (RNP) complexes targeting exon 2 to generate a *RASGRP1*-KO clone (RG1-KO) with loss of RasGRP1 protein (Fig 3A, 3-Suppl A, B). An unedited clone (unedited-i) was used as control. Having established this platform, we targeted autoimmunity-associated enhancer 1 (Fig 3, Suppl A). Analysis of selected single cell clones revealed two clones with deletions in different regions (Enh-i and Enh-ii) that we selected as well a an unedited clone (unedited-ii) as additional control (Figs 3B, 3-Suppl A, B). qPCR analysis revealed reduced levels of *RASGRP1* mRNA expression with a concomitant reduction of RasGRP1 protein levels in clone Enh-ii (Figs 3C, D).

**Figure 3.**
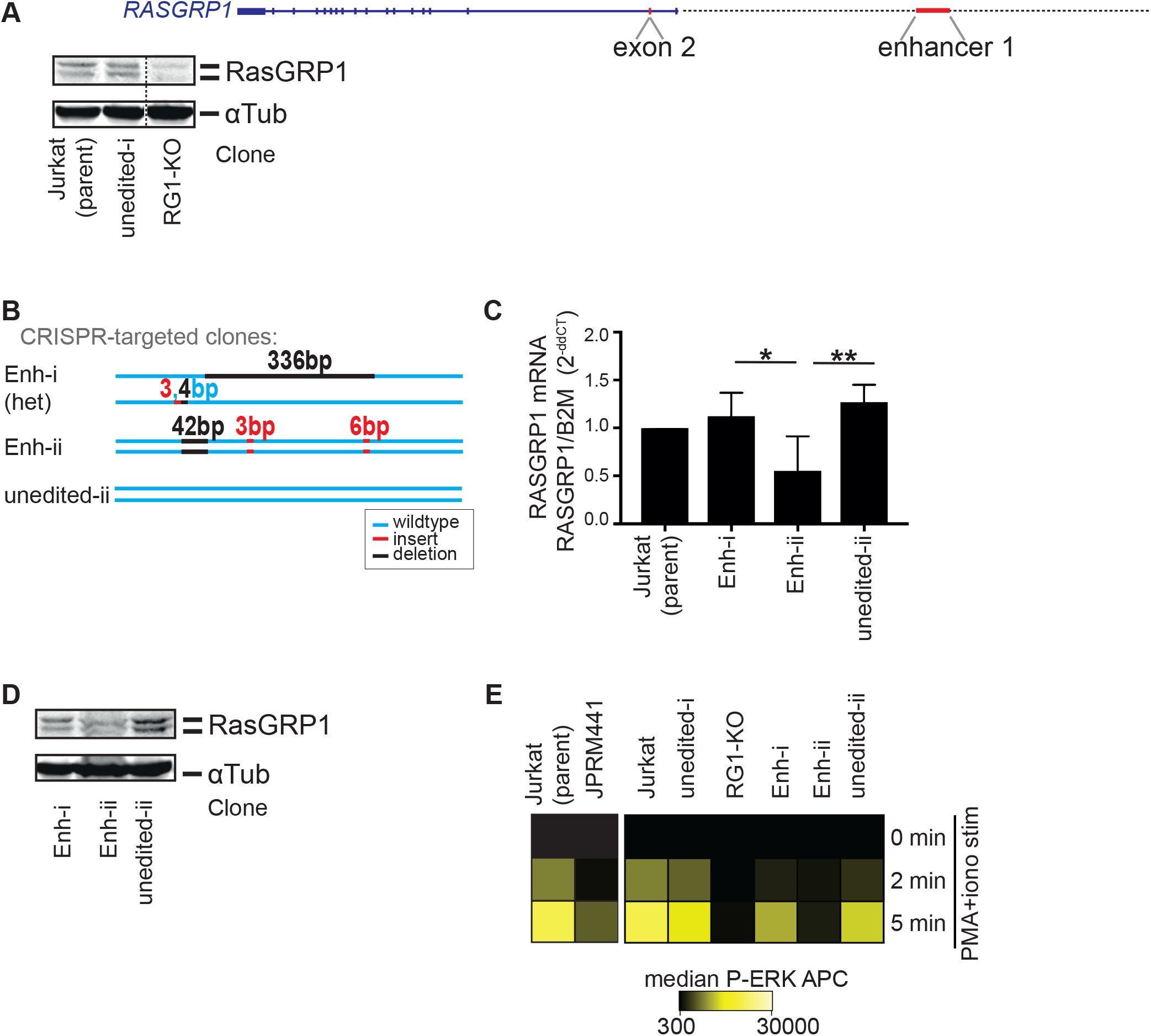
Autoimmunity-associated enhancer 1 regulates *RASGRP1* expression. (A) CRISPR-Cas9 RNP editing of exon 2 in Jurkat cells. Western blot showing RasGRP1 and a-tubulin, parental Jurkat cells, one non-edited clone (unedited-i), and one *RASGRP1*-KO clone (RasGRP1-KO). (B) CRISPR-Cas9 targeting of autoimmunity-associated enhancer 1 in Jurkat cells. Displayed are sequences of two mutants and one unedited clone, showing the WT sequence (blue), deletions (black), and insertions (red), Het= heterozygous. (C-D) RASGRP1 expression levels in parental Jurkat cells, unedited controls, and Enh1 targeted clones. (C) RASGRP1 mRNA expression (RASGRP1/B2M, relative to parent Jurkat cells: 2^−ddCT^). Shown are 4 samples averages ± S.D. 2 separate timepoints of RNA extractions, and duplicates were run per experiment. One-way ANOVA was performed, significance is indicated, * p<0.05, **<0.01. (D) RASGRP1 western blot.(E) Phospho-flow P-ERK analysis of clones at baseline, and after 2 or 5 minutes of PMA/ionomycin stimulation. Shown is a heatmap of median P-ERK expression of one representative experiment out of 3. JPRM441 is a Jurkat line with low (10%) RasGRP1 expression.

Next, we capitalized on a previously optimized platform to analyze RasGRP1 signaling through flow cytometry analyses of phosphorylated ERK a kinase in the canonical Ras-MAPK pathway [2] (Phospho-flow). Phospho-flow allows quantitative detection of defects in ERK signaling such as in JPRM441, a Jurkat line expressing low levels of endogenous RasGRP1 [25]. Phospho-flow analyses revealed that parental Jurkat cells as well as the unedited control clones displayed robust induction of phosphorylated ERK (P-ERK) levels upon stimulation (Figs 3E, 3-Suppl C, D) and impaired induction of P-ERK in JPRM441, as previously reported [25]. Notably, both the *RASGRP1*-knockout clone (RG1-KO) and clone Enh-ii demonstrated impaired induction of P-ERK (Figs 3E, 3-Suppl C, D).

We aimed to mechanistically understand how this enhancer region is impacting *RasGRP1* expression and we postulated that factors binding to the *RasGRP1* enhancer may be distinct in clones Enh-i and Enh-ii. To identify transcription factors regulating *RASGRP1* expression by binding autoimmunity-associated enhancer 1, we performed AP-MS (affinity purification mass spectrometry) analysis [26]. The scrambled sequence of the 42 base pairs (Fig 4 Suppl A) that were deleted in clone Enh-ii, resulted in a loss of binding of CBFB and RUNX1, which together form a transcriptional heterodimer, as well as loss of binding of several ZBTB family members (Fig 4A). We focused on RUNX1 because this factor has been identified as an important transcriptional regulator in hematopoiesis[27] and T cell development[28], during T cell responses[29], and mutations in RUNX1 have been described to play a role in T-ALL [30]. Furthermore, variants in RUNX1 have been associated with autoimmunity[31, 32], and RUNX1 binding sites have been associated with super-enhancers in JIA[33].

**Figure 4.**
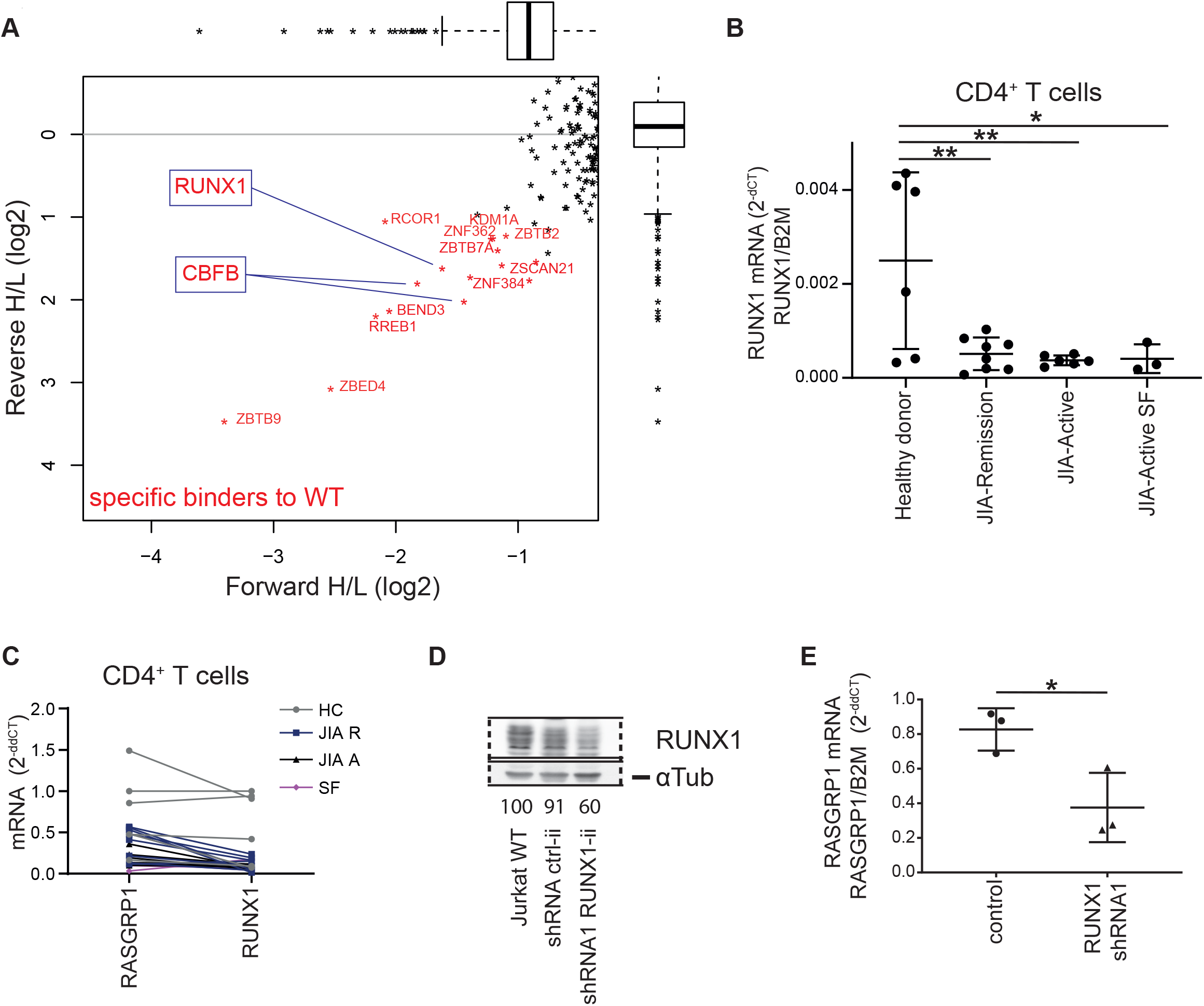
Transcription factor RUNX1 drives RASGRP1 expression, and is reduced in autoimmunity. (A) Affinity-purification mass spectrometry analysis of proteins from jurkat lysates binding wildtype (WT) versus mutant (42 bp scramble) enhancer sequence. In red are factors preferentially bound to WT. (B,C) RUNX1 mRNA levels in CD4+ T cells isolated from juvenile idiopathic arthritis patients during disease remission blood (JIA-R(emission, N=8), active inflammatory disease blood (JIA-A(ctive), N=6), and synovial fluid (JIA-Active SF, N=3), and healthy adult donors blood (N=6). Shown are averages ± S.D. 2^−dCT^ values of RUNX1 corrected for B2M. (C) Plot comparing 2^−ddCT^ RASGRP1 and RUNX1 mRNA expression in each sample, indicated with a line. (D) Western blot of a representative example of RUNX1 knockdown upon RUNX1 shRNA transduction in Jurkat cells. (E) RASGRP1 mRNA, corrected for B2M expression in Jurkat, comparing cells transduced with RUNX1 targeting shRNA or non-targeting (control) shRNA, relative to untransduced Jurkat cells (2-ddCT). Shown are averages ± S.D. One-way anova was performed in B (Holm-Sidak’s post test), two-tailed T-test was performed in E, significance is indicated, * p<0.05, **p<0.01.

To demonstrate the relevance of RUNX1 in CD4^+^ T cells in autoimmunity, we analyzed *RUNX1* expression levels in patients with JIA. We show a decreased *RUNX1* expression in patients with active disease, which again was most pronounced in the synovial fluid, similar to *RASGRP1* expression (Figs 4 B,C). Upon short hairpin RNA-mediated knockdown of *RUNX1* in Jurkat cells (35% knockdown, Figs 4D, Fig 4-Suppl B, C), a clear reduction of *RASGRP1* mRNA expression occurred (Figs 4E). These data provide a novel link between RUNX1 and RasGRP1, and a mechanistic explanation for the reduced RasGRP1 expression in clone Enh-ii, because RUNX1 can no longer bind to the regulatory enhancer element that is key for *RASGRP1* transcription.

## Discussion

We show for the first time that decreased expression levels of Rasgrp1 induce autoimmunity in mice through a reduction of positive thymocyte selection, and that decreased RASGRP1 in CD4^+^ T cells is specifically detected under circumstances of active autoimmunity in patients. In mice, reduced expression of RASGRP1 mildly disturbs T cell development, and results in a delayed onset of disease compared to complete deletion of RASGRP1. We detected reduced levels of RASGRP1 in peripheral CD4^+^ T cells in patients, specifically in the synovial fluid at the site of active inflammation, suggesting that dysregulation of RASGRP1 expression occurs in activated T cells. In line with this, we identified a *RASGRP1* enhancer containing a cluster of autoimmunity-risk SNPs, that showed a higher H3K27ac signal in previously activated CD4^+^ memory T cells.

Mechanistically, we show that transcription factor RUNX1 is essential for regulation of RASGRP1 expression levels. RUNX1 has been identified as an important transcription factor in hematopoiesis and T cell development, and mutations in RUNX1 have been described to play a role in T-ALL [30]. To date, RUNX1 has not been linked to RASGRP1 transcriptional regulation, and the role of RUNX1 in autoimmunity remained ambiguous. For example, in patients with active immune thrombocytopenia RUNX1 mRNA expression was elevated [34], and RUNX1 has been suggested to promote maturation of CD4^+^ T cells in mice [35]. By contrast, another study showed that RUNX1 deletion in mice led to autoimmune lung disease [36]. Here we provide evidence that decreased expression of RUNX1 leads to reduced RASGRP1 transcription, suggesting a role for RUNX1 in fine-tuning Ras-MAPK signaling in both developing T cells, and peripheral CD4^+^ T cells.

In conclusion, RasGRP1 expression regulates TCR-induced signaling and thymocyte selection in a dose-dependent manner. Tight regulation of RasGRP1 expression is vital to maintain immunological health and prevent diseases such as leukemia, autoimmunity and immune deficiency (Fig. 5).

**Figure 5.**
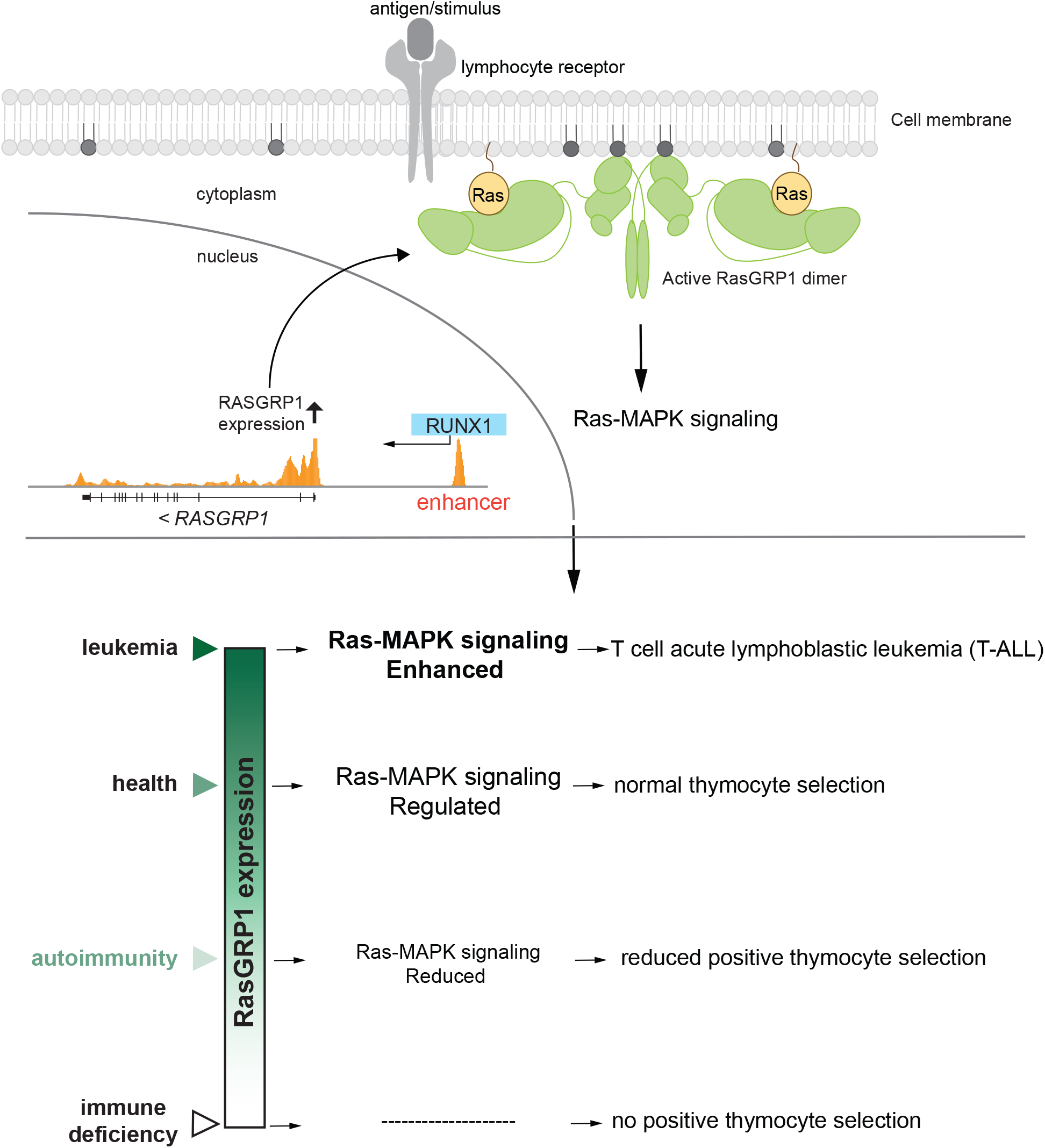
Tight regulation of RASGRP1 expression in T cells is vital to prevent disease. ***Top:*** RASGRP1 transcription is regulated by enhancer 1 binding transcription factors (RUNX1/CBFB). T cell receptor signaling upon antigen/MHC binding induces activation of the RasGRP1 dimer, allowing membrane recruitment and binding to Ras. Next, RasGRP1 releases GDP bound to Ras, allowing GTP binding and active Ras-MAPK signaling. ***Bottom:*** The level of RASGRP1 expression and resulting Ras-MAPK signaling affects thymocyte selection and clinical outcome: Reduced RASGRP1 expression results in lower Ras-MAPK signaling, thus reduced positive thymocyte selection signals, and can cause autoimmunity. Absence of RASGRP1 results in a complete loss of positive thymocyte selection and immune deficiency, while increased levels of RASGRP1 expression and Ras-MAPK signaling have been shown in T-ALL. Together, tight regulation of RasGRP1 expression is essential to maintain healthy T cells and prevent disease.

## Materials and methods

### Patients and healthy donors

To investigate the differences in expression levels during disease course, patients with oligoarticular JIA (oJIA) were selected of which at least 30*106 PBMCs from both active disease as well as remission were available (N=8, Table 1). In addition, three unpaired samples of oJIA SFMCs were selected. Healthy adult volunteers (n=6) derived PBMCs, collected via the UMCU facility, served as a control group.

### Mice

C57BL/6 mice wildtype, Rasgrp1^−/−^ (provided by J. Stone), and Rasgrp1^+/−^ were bred in house at UCSF.

### Study approval

This study was conducted according to the principles expressed in the Declaration of Helsinki, and was approved by the UMCU medical ethical committee, study Pharmachild, number 11-499/C.

All donors provided written informed consent prior to inclusion in this study for sample collection and analysis, and all donors were numbered to anonymize the samples.

Mice were housed and treated in accordance with the guidelines of the Institutional Animal Care and Use Committee (IACUC) guidelines of the University of California, San Francisco (protocol number AN098375-03B).

### HEp-2 ANA Assays

HEp-2 assays were performed utilizing the Nova-Lite kit from INOVA diagnostics. See supplementary methods.

### Cells, medium and reagents

PBMCs and SFMCs were extracted using Ficoll-Paque PLUS density gradient centrifugation (GE Healthcare Life Sciences), and stored frozen in FCS+10% DMSO. Jurkat and JPRM441 (low RasGRP1) cells were previously characterized and described [25, 37]. RPMI1640 containing 10 mM HEPES, 2mM L-glutamine, 100 U/ml Penicillin-streptomycin, and 10% FCS was used as culture medium. Starvation was performed in plain RPMI1640 for 30 minutes. When indicated cells were stimulated with PMA (sigma, 20ng/ml), ionomycin (1 uM).

### Flow cytometry

FACS buffer consisted of Phosphate buffered salt (PBS) with 2 mM EDTA, 2% FCS, and 0.1% NaN3. Cells were harvested, blocked with Fc block (1:200), normal mouse serum, normal rat serum, next cells were labeled with viability dye (eBiosciences eF780 or eF506 fixable viability dye, 1:1000), and extracellular staining was performed by incubation of cells with antibodies in FACS buffer for 15 minutes on ice.

#### Phospho-flows

cells were washed, resuspended in plain RPMI, seeded 0.4 × 10^6^ per well in a 96 well round bottom plate, and starved in an incubator for 30 min. After stimulation, cells were fixated in prewarmed fixations buffer (Cytofix, BD biosciences), or in 2 % paraformaldehyde in Phosphate buffered saline (PBS), for 15 min at RT, washed in FACS buffer, and permeabilized in MetOH for 30 min on ice. After washing with FACS buffer, cells were incubated with anti-phospho-ERK, washed, and incubated with secondary antibody. Cells were then washed and run on a BD FACSCanto.

### Magnetic cell isolation of CD4^+^ T cells

CD4^+^ T cells were magnetically purified using CD4^+^ T cell isolation kit and the autoMACS® Columns Pro Separator machine (Miltenyi Biotec), according to the manufacturer instructions. Purity was assessed by flow cytometry staining, and manual gating in FacsDIVA (BD Biosciences).

### Antibodies

#### MACS purity

CD3-BV510, clone OKT3, Biolegend; CD4-FITC, clone RPA-T4, eBioscience.

#### Phospho-flow

P-ERK (clone 197G2, #4377 s, diluted 1:50). AffiniPure F(ab’)2 fragment Donkey-anti-Rabbit IgG, conjugated to APC (#711-136-152, 1:50) or PE (#711-116-152, 1:50, Jackson ImmunoResearch,).

#### Antibodies for analyzing thymocyte development

BD Pharmingen PerCP-Cy5.5 anti-mouse CD4 (clone RM4-5; cat# 550954) (dil 1:800)

UCSF mAb core FITC anti-mouse CD8 (clone YTS169) (dil 1:800)

BD Pharmingen™ PE Anti-mouse CD5 (clone 53-7.3; cat# 553023) (dil 1:100)

Tonbo APC anti-mouse CD25 (clone PC61.5; cat# 20-0251) (dil 1:100)

Biolegend PE/Cy7 anti-mouse TCR- β (clone H57-597; cat# 109221) (dil 1:200)

Tonbo PE/Cy7 anti-human/mouse CD44 (clone IM7; cat# 60-0441-U025) (dil 1:100)

BD Pharmingen APC anti-mouse CD69 (clone H1.2F3; cat# 560689) (dil 1:100)

#### Dump channel abs (violetFluor 450) (dil 1:100)

Tonbo Biosciences violetFluor 450 anti-human/mouse CD45R/CD19/b220 (clone RA3-6B2; cat# 75-0452-U100) ThermoFisher eFluor 450 anti-mouse Ter-119 (cat# 48-5921-82), Tonbo Biosciences violetFluor 450 anti-human/mouse CD11b (clone M1/70; cat# 75-0112-U100),Tonbo Biosciences violetFluor 450 anti-human/mouse CD11c (clone N418; cat# 75-0114-U025), Tonbo Biosciences violetFluor 450 anti-mouse Gr-1 (clone RB6-8C5; cat# 75-5931-U025).

#### Viability dye

Live/dead fixable violet dead cell stain kit for 405 nm excitation, ThermoFisher Scientific (cat# L34955) (dil 1:1000)

### HEp-2 ANA Assays

HEp-2 assays were performed utilizing the Nova-Lite kit from INOVA diagnostics. Serum was applied to slides, stained with IgG-FITC (Jackson Labs) and DAPI (500ng/ml, LifeTechnologies). Slides were imaged on a Keyence BZ-X710 microscope.

Sera were scored blindly by 3 separate researchers as ANA negative or positive. Each slide contained a no serum negative control or a CD45 Wedge B6-129 F1 positive control serum (a gift from the Hermiston lab, UCSF).

### CRISPR-Cas9 editing

#### Sequences of crRNA

exon 2: GTCAATGAGATCGTCCAGGC, and AGCTGTCAATGAGATCGTCC, Enhancer: TTATAAGAAGGGCTTACCGTGGG, TTTATAAGAAGGGCTTACCGTGG, CAAAACGGAGTTACATAGCAAGG, TGCTTGATCTCAGATTAAGCAGG, CTTAATCTGAGATCAAGCACAGG.

### DNA and RNA

DNA and RNA were extracted from cells using either Blood & Cell Culture DNA Mini Kit and RNAeasy isolation kit (Qiagen), or all-prep DNA/RNA kit (Qiagen). To assess genome editing, PCR amplification was followed by PCR-clean up and send out for sequencing, using the following primers: Exon 2: Fwd 5’ GAAACCTTCCCATGGCTGCA 3’, Rev 5’ TGCAGCTGTCAATGAGATCGT 3’; Enhancer: Fwd 5’ GGATGGGCTGGTTGAGTCAA 3’, Rev 5’ ACAGTGTAGGTTCCTAGACCCT 3’. PCR fragments were sequenced and analyzed using TIDE [38]. Subsequently PCR fragments were cloned into pJET (Thermo Scientific) and sequenced with primers from the pJET kit.

RNA purity and concentration were measured by NanoDrop 2000 (Thermo Scientific). cDNA was synthesized per 15 ng total RNA using the iScriptTM cDNA Synthesis Kit (Bio-Rad). Quantitative amplification of the cDNA templates was assessed by the SYBR® Select Master Mix (Thermo-Scientific) using the QuantStudio 12K Flex (Thermo-Scientific), or CFX384 Touch (Bio-Rad).

Samples contained 4 ng cDNA with primers for β-2-microglobulin (forward, 5’ TGCTGTCTCCATGTTTGATGTATCT 3’; reverse 5’ TCTCTGCTCCCCACCTCTAAGT 3’), RUNX1 (forward, 5’ TGAGTCATTTCCTTCGTACC 3’; reverse 5’ TGCTGGCATCGTGGA 3’), or RasGRP1 (PrimePCR SYBR Green Assay RasGRP1 human, Bio-Rad, amplicon context sequence GGTTCCTTGGTTCCCGGGCATAGGAAAGCTCATAGATTTCATCCTCAGTGTAGTAAAGATCCAGGGATAACGTCAGCAAGTGTACCAAGTCCTTGTTAGCCTCCAAGG (exon 10). Each run included H2O as negative control, and each sample was run in triplicate.

### DNA affinity purification and LC-MS analysis

Nuclear extracts from Jurkat T cells were generated as described [39]. Oligonucleotides for the DNA affinity purifications were ordered from IDT with the forward strand containing a 5’ biotin moiety. DNA affinity purifications and on-bead trypsin digestion was performed as described [26]. Tryptic peptides from SNP variant pull-downs were desalted using Stage (stop and go extraction) tips [40], and then subjected to stable isotope di-methyl labeling [41] on the Stage tips. Matching light and heavy peptides were then combined and samples were finally subjected to LC-MS and subsequent data analyses using MaxQuant [42] and R essentially as described.

### shRNA transduction

Lentiviral particles were produced in HEK293T cells using third generation packaging factors. shRNA plasmids were used from the Mission library (Sigma), for RUNX1 (TRCN0000338427, TRCN0000338490, and TRCN0000338489), and non-targeting shRNA control (SHC002). Jurkat cells were infected by spinoculation in the presence of 8μg/ml of polybrene for 3 hours at 22°C at 800rpm in a table-top centrifuge. Cells were selected with 1μg/ml of puromycin 24 hours after infection.

### Western Blot

Cells were lysed in SDS buffer, and lysates were run on SDS-page gel and transferred to a PVDF membrane. After blocking for 1 hour in 3% Bovine Serum Albumin (Roche) in TBS-T, membranes were incubated overnight at 4°C with primary antibodies for RasGRP1 (JR-E80-2[9], 1:1500), RUNX1 (ab23980, Abcam, 1:1000), or α-tubulin (T6074, Sigma, 1:2000) in blocking buffer, followed by secondary antibodies in TBS-T for 1 hour at RT. Proteins were visualized using enhanced chemiluminescence.

## Author contributions

MB, TD, DS, DM, SB, KK, SZ, YV, conducted experiment, acquired and analyzed data. FvW, SdR, and AM provided resources and designed experiments. JR and YV designed the study, and wrote the first draft. All authors revised the manuscript.

## Acknowledgements

The authors wish to thank Oghenekevwe Michelle Gbenedio, Flow Cytometry Core (NIH P30DK063720, UCSF), Marten Hornsveld (LUMC), Fried Zwartkruis, Jose Ramos-Pittol, Rina Wichers, Flow Cytometry Core (UMCU) for assistance.

Funding was provided by cancergenomicscenter.nl, and Marie Curie PIOF-GA-2012-328666 (YV), NSF-GRFP (1650113 to DRM) and NIH-NIAID (R01-AI104789 and P01-AI091580 to JPR), Jeffrey G. Klein Family Fellowship in Diabetes (DRS), NIH-NIAID (DP3DK111914-01 (NIDDK) and R01DK1199979 (NIDDK), to AM), Innovative Genomics Institute (IGI, to AM). A.M. holds a Career Award for Medical Scientists from the Burroughs Wellcome Fund, is an investigator at the Chan Zuckerberg Biohub, member of the Parker Institute for Cancer Immunotherapy (PICI).

The Vermeulen lab is in the Oncode Institute, which is co-funded by the Dutch Cancer Society (KWF).

## Conflicts of interest disclosure

Jeroen Roose is a co-founder and scientific advisor of Seal Biosciences, Inc. and on the scientific advisory committee for the Mark Foundation for Cancer research. A.M. is a co-founder of Arsenal Biosciences, Spotlight Therapeutics and Trizell, serves as on the scientific advisory board of PACT Pharma and was a former advisor to Juno Therapeutics. The Marson Laboratory has received sponsored research support from Juno Therapeutics, Epinomics, Sanofi and a gift from Gilead. D.R.S. is a co-founder of Beeline Therapeutics. The authors have no additional financial interests.

## Figure supplement legends

**Figure 1-Supplement.**
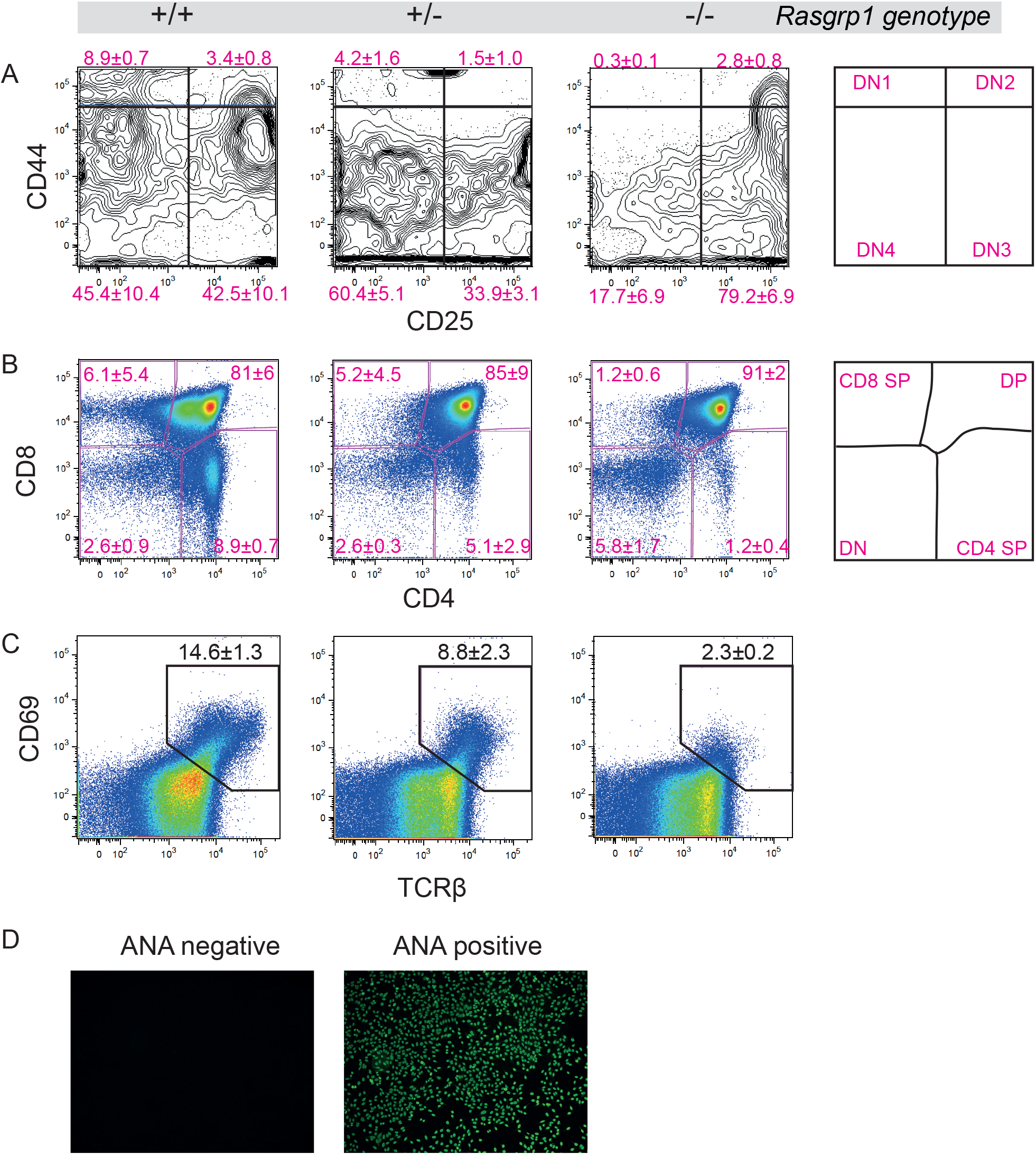
Limited RasGRP1 expression results in dose-dependent autoimmunity in mice. (A-C) Representative dotplots of flow cytometry analysis of thymocytes (at 5 weeks) from RASGRP1 wildtype (+/+), heterozygous (+/−), and KO (−/−) mice. (A-C) Live, singlet gated thymocytes. Indicated are average percentages ± S.D. (A) Total thymocytes, CD4CD8 DN, SP, DP populations. (B) CD44^−^CD25^+^ DN (DN3) thymocytes. (C) TCRß^+^CD69^+^ DP thymocytes. (D) Representative examples of ANA stainings, 1 negative staining (left), one positive staining (right).

**Figure 2-Supplement A.**
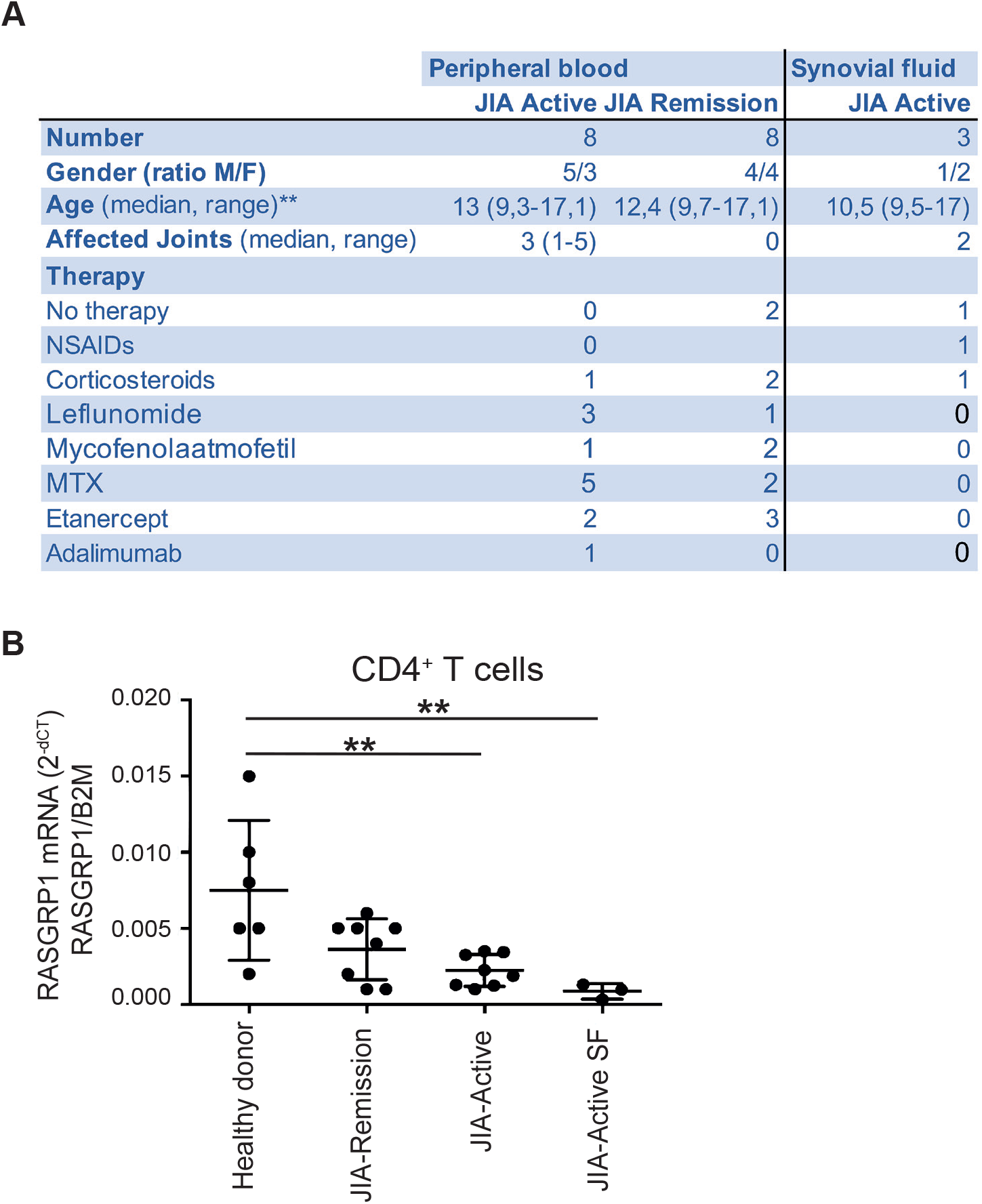
RASGRP1 levels and expression regulation are associated with human autoimmunity. (A) Characteristics of included JIA patients. (B) RASGRP1 mRNA levels in CD4+ T cells were isolated from juvenile idiopathic arthritis patients during disease remission blood (JIA-Remission, N=8), active inflammatory disease blood (JIA-Active, N=8), and synovial fluid (JIA-Active SF, N=3), and healthy adult donors blood (N=6). Shown are averages ± S.D. 2^−dCT^ values of RasGRP1 corrected for B2M. One-way Anova, with Holm-Sidak’s multiple comparisons test was used for statistical analysis.

**Figure 2-Supplement B.**
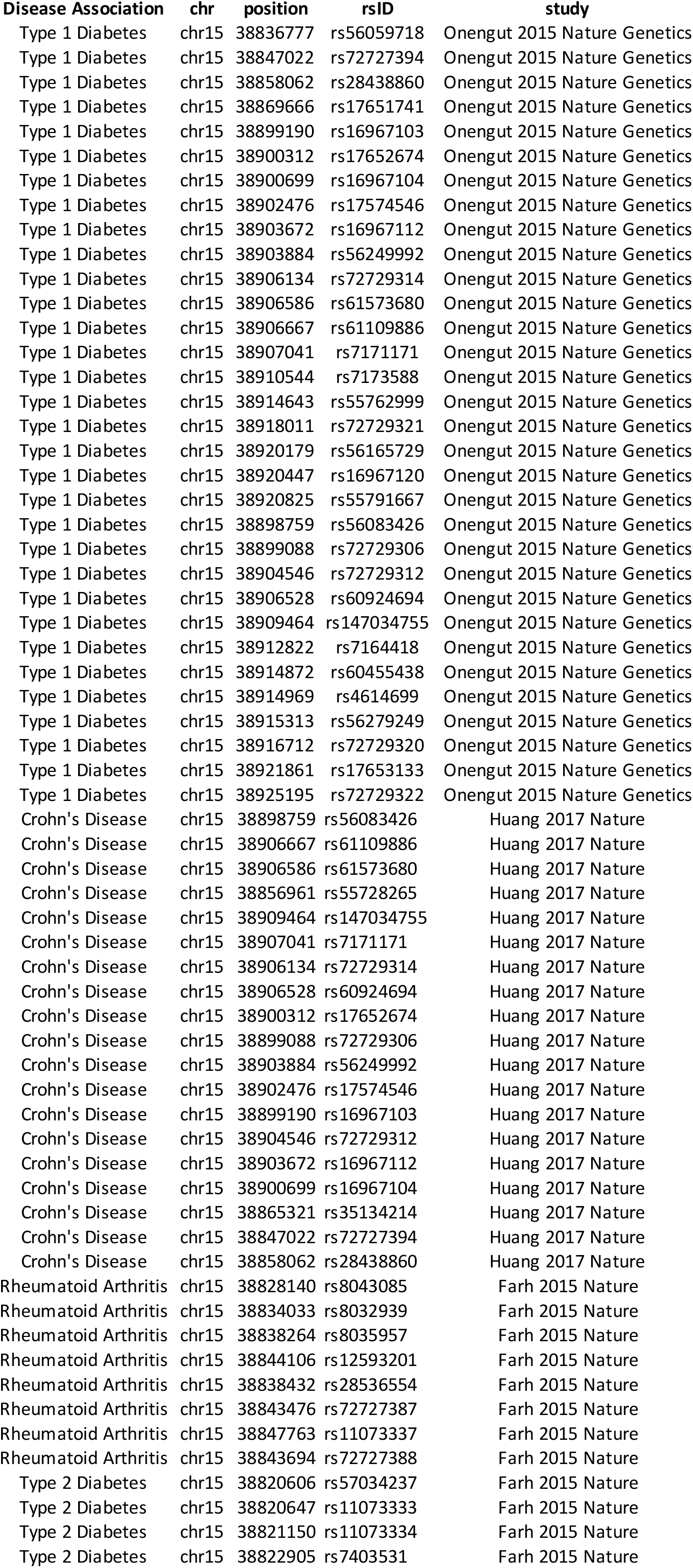
Rasgrp1 SNPs associated with autoimmunity

**Figure 3-Supplement.**
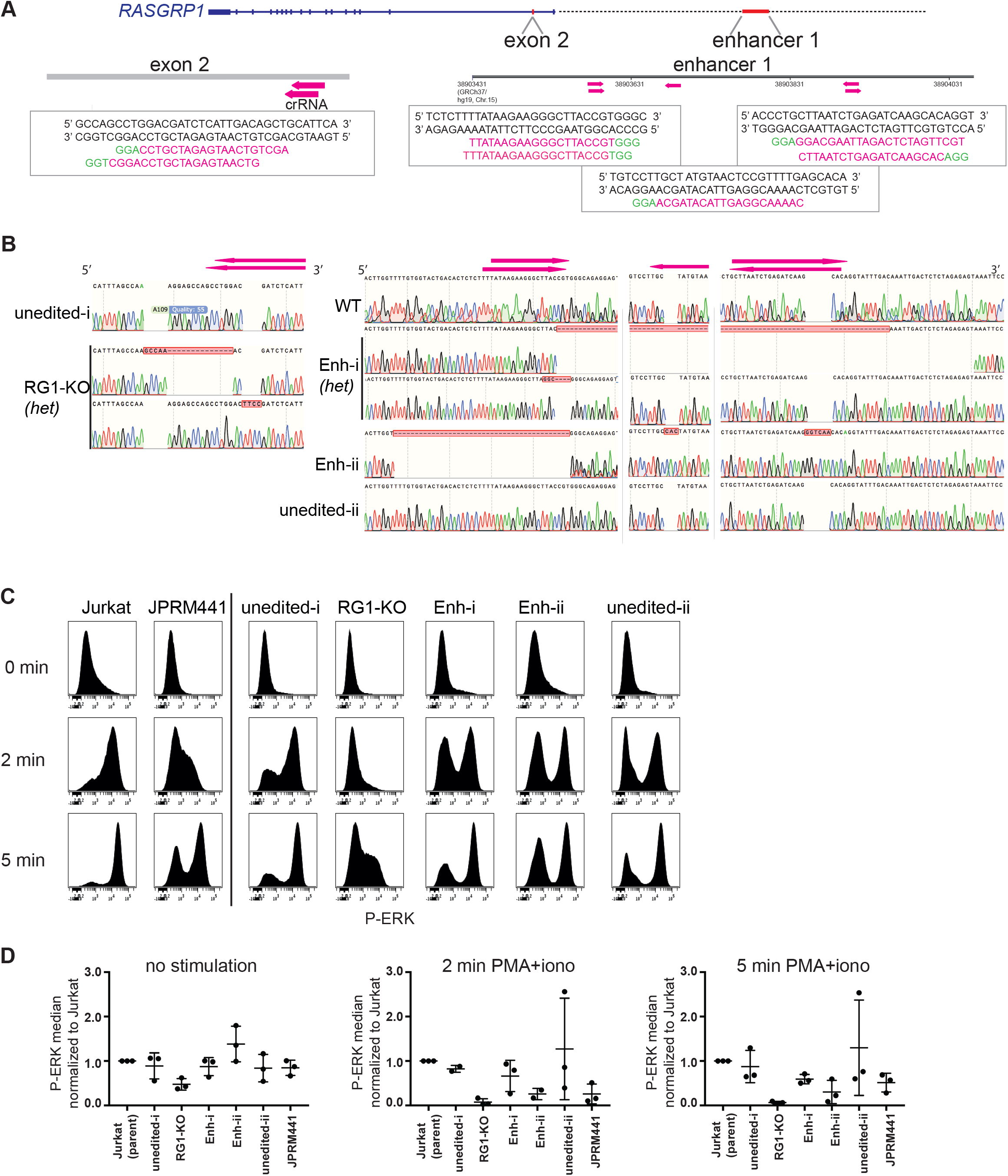
Autoimmunity-associated enhancer 1 regulates RASGRP1 expression. A) Enhancer 1 (autoimmunity hotspot 1) was targeted with CRISPR-Cas9 RNP editing in Jurkat cells. As a control we targeted exon 2 to generate *RASGRP1*-KO clones. Indicated are the crRNA used (pink arrows and sequences, PAM in green). (B) Sequencing profiles of clones from Snapgene alignments, deletions and inserts are marked in red boxes, pink arrows indicate crRNA. Left: exon 2 targeted clone showing complete RasGRP1 protein loss, mutations. Right: Enhancer 1-targeted clones, enh1-ii (42bp del), unedited-ii are homozygous, enh1-i is heterozygous, with one 336 bp deletion, and one indel (3 bp ins, 4 bp del). (C,D) Phospho-flow analysis of P-ERK at baseline (0 min), or upon stimulation with PMA/ionomycin (2 min, 5 min). (C) Representative histograms. (D) Average of separate experiments showing median P-ERK normalized to parent Jurkat. (N=3, only 2 min PMA+iono Enh1-ii: N=2).

**Figure 4-supplement.**
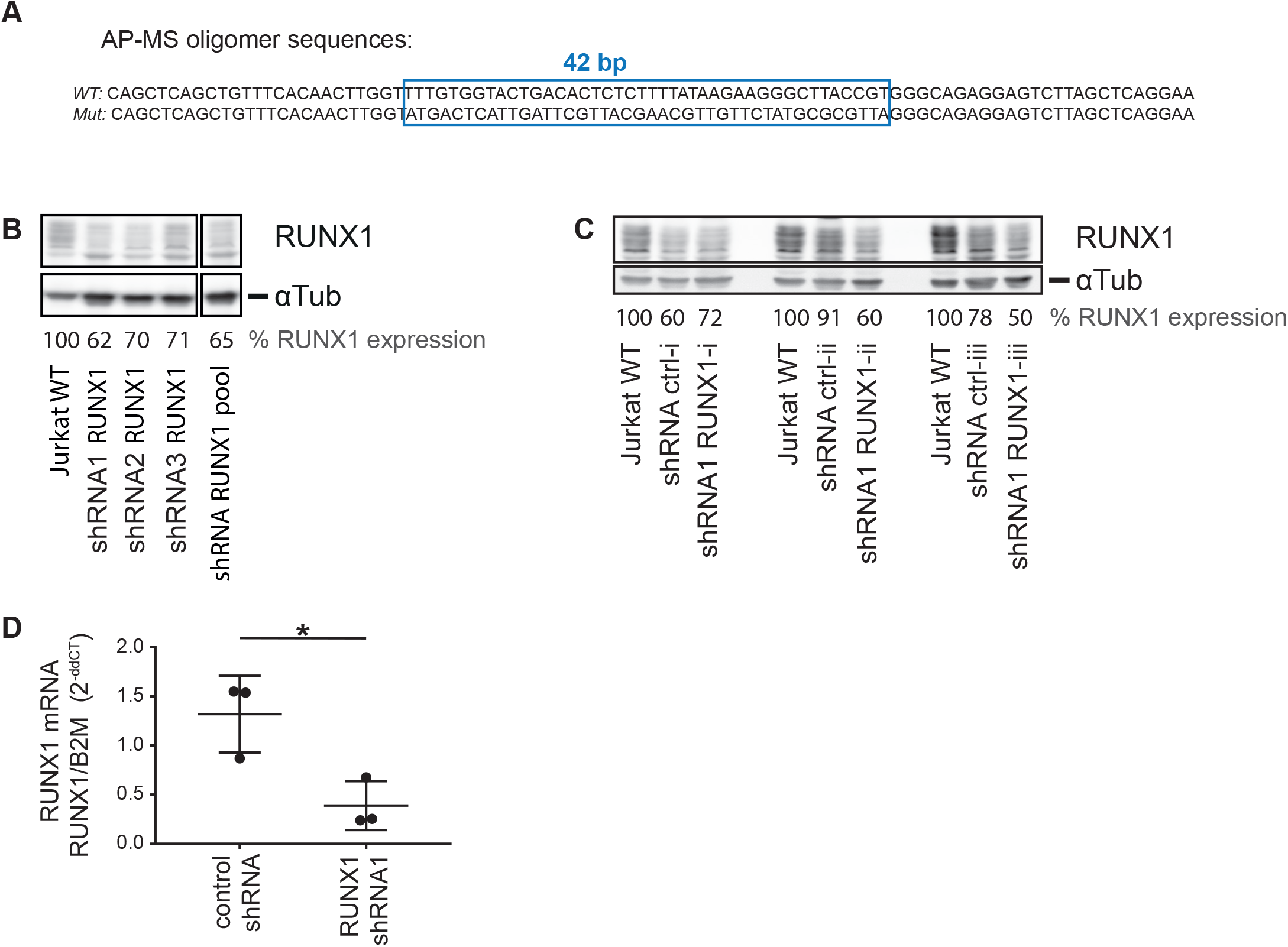
Transcription factor RUNX1 drives RASGRP1 expression, and is reduced in autoimmunity. (A) Sequences of oligomers used for AP-MS. (B,C) Western blot of Jurkat lysates 48hrs after upon shRNA transduction, displaying expression of RUNX1 and a-Tubulin, to assess knockdown. RUNX1 protein expression was normalized to untransduced Jurkat cells. (B) Jurkat cells were transduced with a pool of 3 shRNA constructs, or single constructs, targeting RUNX1. (C) Biological replicates of Jurkat cells transduced with non-targeting control, or RUNX1 targeting shRNA1. (D) RUNX1 mRNA expression 24 hrs after transduction with non-targeting control, or RUNX1 targeting shRNA, relative to untransduced Jurkat.

## Notes

#### Summary of Updates

We added novel data figures (figure 4), showing that RUNX1 expression is low in CD4+ T cells from patients with autoimmunity, similar to RASGRP1, and that RUNX1 drives RASGRP1 expression.

## References

[1] O. Ksionda, A. Limnander, J. P. Roose. RasGRP Ras guanine nucleotide exchange factors in cancer. Front Biol (Beijing), 2013;8:508–32.

[2] Y. Vercoulen, Y. Kondo, J. S. Iwig, A. B. Janssen, K. A. White, M. Amini et al. A Histidine pH sensor regulates activation of the Ras-specific guanine nucleotide exchange factor RasGRP1. Elife, 2017;6.

[3] J. S. Iwig, Y. Vercoulen, R. Das, T. Barros, A. Limnander, Y. Che et al. Structural analysis of autoinhibition in the Ras-specific exchange factor RasGRP1. Elife, 2013;2:e00813.

[4] N. A. Dower, S. L. Stang, D. A. Bottorff, J. O. Ebinu, P. Dickie, H. L. Ostergaard et al. RasGRP is essential for mouse thymocyte differentiation and TCR signaling. Nat Immunol, 2000;1:317–21.

[5] J. J. Priatel, S. J. Teh, N. A. Dower, J. C. Stone, H. S. Teh. RasGRP1 transduces low-grade TCR signals which are critical for T cell development, homeostasis, and differentiation. Immunity, 2002;17:617–27.

[6] E. Salzer, D. Cagdas, M. Hons, E. M. Mace, W. Garncarz, O. Y. Petronczki et al. RASGRP1 deficiency causes immunodeficiency with impaired cytoskeletal dynamics. Nat Immunol, 2016;17:1352–60.

[7] S. Winter, E. Martin, D. Boutboul, C. Lenoir, S. Boudjemaa, A. Petit et al. Loss of RASGRP1 in humans impairs T-cell expansion leading to Epstein-Barr virus susceptibility. EMBO Mol Med, 2018;10:188–99.

[8] H. Mao, W. Yang, S. Latour, J. Yang, S. Winter, J. Zheng et al. RASGRP1 mutation in autoimmune lymphoproliferative syndrome-like disease. J Allergy Clin Immunol, 2017.

[9] C. Hartzell, O. Ksionda, E. Lemmens, K. Coakley, M. Yang, M. Dail et al. Dysregulated RasGRP1 responds to cytokine receptor input in T cell leukemogenesis. Sci Signal, 2013;6:ra21.

[10] O. Ksionda, A. A. Melton, J. Bache, M. Tenhagen, J. Bakker, R. Harvey et al. RasGRP1 overexpression in T-ALL increases basal nucleotide exchange on Ras rendering the Ras/PI3K/Akt pathway responsive to protumorigenic cytokines. Oncogene, 2015.

[11] S. Yasuda, R. L. Stevens, T. Terada, M. Takeda, T. Hashimoto, J. Fukae et al. Defective expression of Ras guanyl nucleotide-releasing protein 1 in a subset of patients with systemic lupus erythematosus. J Immunol, 2007;179:4890–900.

[12] M. L. Golinski, T. Vandhuick, C. Derambure, M. Freret, M. Lecuyer, C. Guillou et al. Dysregulation of RasGRP1 in rheumatoid arthritis and modulation of RasGRP3 as a biomarker of TNFalpha inhibitors. Arthritis Res Ther, 2015;17:382.

[13] K. K. Farh, A. Marson, J. Zhu, M. Kleinewietfeld, W. J. Housley, S. Beik et al. Genetic and epigenetic fine mapping of causal autoimmune disease variants. Nature, 2015;518:337–43.

[14] H. Huang, M. Fang, L. Jostins, M. Umicevic Mirkov, G. Boucher, C. A. Anderson et al. Fine-mapping inflammatory bowel disease loci to single-variant resolution. Nature, 2017;547:173–8.

[15] S. Onengut-Gumuscu, W. M. Chen, O. Burren, N. J. Cooper, A. R. Quinlan, J. C. Mychaleckyj et al. Fine mapping of type 1 diabetes susceptibility loci and evidence for colocalization of causal variants with lymphoid gene enhancers. Nat Genet, 2015;47:381–6.

[16] S. R. Daley, K. M. Coakley, D. Y. Hu, K. L. Randall, C. N. Jenne, A. Limnander et al. Rasgrp1 mutation increases naive T-cell CD44 expression and drives mTOR-dependent accumulation of Helios+ T cells and autoantibodies. Elife, 2013;2:e01020.

[17] R. S. Sellers, C. B. Clifford, P. M. Treuting, C. Brayton. Immunological variation between inbred laboratory mouse strains: points to consider in phenotyping genetically immunomodified mice. Vet Pathol, 2012;49:32–43.

[18] D. P. Golec, N. A. Dower, J. C. Stone, T. A. Baldwin. RasGRP1, but not RasGRP3, is required for efficient thymic beta-selection and ERK activation downstream of CXCR4. PLoS One, 2013;8:e53300.

[19] M. J. Bevan, K. A. Hogquist, S. C. Jameson. Selecting the T cell receptor repertoire. Science, 1994;264:796–7.

[20] J. J. Coughlin, S. L. Stang, N. A. Dower, J. C. Stone. RasGRP1 and RasGRP3 regulate B cell proliferation by facilitating B cell receptor-Ras signaling. J Immunol, 2005;175:7179–84.

[21] J. C. Stone. Regulation and Function of the RasGRP Family of Ras Activators in Blood Cells. Genes Cancer, 2011;2:320–34.

[22] D. Hnisz, B. J. Abraham, T. I. Lee, A. Lau, V. Saint-Andre, A. A. Sigova et al. Super-enhancers in the control of cell identity and disease. Cell, 2013;155:934–47.

[23] S. Pott, J. D. Lieb. What are super-enhancers? Nat Genet, 2015;47:8–12.

[24] K. Schumann, S. Lin, E. Boyer, D. R. Simeonov, M. Subramaniam, R. E. Gate et al. Generation of knock-in primary human T cells using Cas9 ribonucleoproteins. Proc Natl Acad Sci U S A, 2015;112:10437–42.

[25] J. P. Roose, M. Mollenauer, V. A. Gupta, J. Stone, A. Weiss. A diacylglycerol-protein kinase C-RasGRP1 pathway directs Ras activation upon antigen receptor stimulation of T cells. Mol Cell Biol, 2005;25:4426–41.

[26] M. M. Makowski, E. Willems, J. Fang, J. Choi, T. Zhang, P. W. Jansen et al. An interaction proteomics survey of transcription factor binding at recurrent TERT promoter mutations. Proteomics, 2016;16:417–26.

[27] T. Okuda, J. van Deursen, S. W. Hiebert, G. Grosveld, J. R. Downing. AML1, the target of multiple chromosomal translocations in human leukemia, is essential for normal fetal liver hematopoiesis. Cell, 1996;84:321–30.

[28] M. Kawazu, T. Asai, M. Ichikawa, G. Yamamoto, T. Saito, S. Goyama et al. Functional domains of Runx1 are differentially required for CD4 repression, TCRbeta expression, and CD4/8 double-negative to CD4/8 double-positive transition in thymocyte development. J Immunol, 2005;174:3526–33.

[29] V. Lazarevic, X. Chen, J. H. Shim, E. S. Hwang, E. Jang, A. N. Bolm et al. T-bet represses T(H)17 differentiation by preventing Runx1-mediated activation of the gene encoding RORgammat. Nat Immunol, 2011;12:96–104.

[30] J. Zhang, L. Ding, L. Holmfeldt, G. Wu, S. L. Heatley, D. Payne-Turner et al. The genetic basis of early T-cell precursor acute lymphoblastic leukaemia. Nature, 2012;481:157–63.

[31] C. Helms, L. Cao, J. G. Krueger, E. M. Wijsman, F. Chamian, D. Gordon et al. A putative RUNX1 binding site variant between SLC9A3R1 and NAT9 is associated with susceptibility to psoriasis. Nat Genet, 2003;35:349–56.

[32] S. Tokuhiro, R. Yamada, X. Chang, A. Suzuki, Y. Kochi, T. Sawada et al. An intronic SNP in a RUNX1 binding site of SLC22A4, encoding an organic cation transporter, is associated with rheumatoid arthritis. Nat Genet, 2003;35:341–8.

[33] J. G. Peeters, S. J. Vervoort, S. C. Tan, G. Mijnheer, S. de Roock, S. J. Vastert et al. Inhibition of Super-Enhancer Activity in Autoinflammatory Site-Derived T Cells Reduces Disease-Associated Gene Expression. Cell Rep, 2015;12:1986–96.

[34] X. Zhong, Y. Wu, Y. Liu, F. Zhu, X. Li, D. Li et al. Increased RUNX1 expression in patients with immune thrombocytopenia. Hum Immunol, 2016;77:687–91.

[35] F. C. Hsu, M. J. Shapiro, B. Dash, C. C. Chen, M. M. Constans, J. Y. Chung et al. An Essential Role for the Transcription Factor Runx1 in T Cell Maturation. Sci Rep, 2016;6:23533.

[36] W. F. Wong, K. Kohu, T. Nagashima, R. Funayama, M. Matsumoto, E. Movahed et al. The artificial loss of Runx1 reduces the expression of quiescence-associated transcription factors in CD4(+) T lymphocytes. Mol Immunol, 2015;68:223–33.

[37] J. P. Roose, M. Mollenauer, M. Ho, T. Kurosaki, A. Weiss. Unusual interplay of two types of Ras activators, RasGRP and SOS, establishes sensitive and robust Ras activation in lymphocytes. Mol Cell Biol, 2007;27:2732–45.

[38] E. K. Brinkman, T. Chen, M. Amendola, B. van Steensel. Easy quantitative assessment of genome editing by sequence trace decomposition. Nucleic Acids Res, 2014;42:e168.

[39] A. H. Smits, P. W. Jansen, I. Poser, A. A. Hyman, M. Vermeulen. Stoichiometry of chromatin-associated protein complexes revealed by label-free quantitative mass spectrometry-based proteomics. Nucleic Acids Res, 2013;41:e28.

[40] J. Rappsilber, M. Mann, Y. Ishihama. Protocol for micro-purification, enrichment, pre-fractionation and storage of peptides for proteomics using StageTips. Nat Protoc, 2007;2:1896–906.

[41] P. J. Boersema, R. Raijmakers, S. Lemeer, S. Mohammed, A. J. Heck. Multiplex peptide stable isotope dimethyl labeling for quantitative proteomics. Nat Protoc, 2009;4:484–94.

[42] J. Cox, M. Mann. MaxQuant enables high peptide identification rates, individualized p.p.b.-range mass accuracies and proteome-wide protein quantification. Nat Biotechnol, 2008;26:1367–72.

[43] C. Roadmap Epigenomics, A. Kundaje, W. Meuleman, J. Ernst, M. Bilenky, A. Yen et al. Integrative analysis of 111 reference human epigenomes. Nature, 2015;518:317–30.

